# Evenness-richness scatter plots: a graphical, intuitive approach to alpha diversity analysis

**DOI:** 10.1101/2020.09.23.310045

**Authors:** Jeff Gauthier, Nicolas Derome

## Abstract

Shannon’s diversity index is a popular alpha diversity metric because it estimates both richness and evenness in a single equation. However, since its value is dependent of both those parameters, there is theoretically an infinite number of richness / evenness value combinations translating into the same index score. By decoupling both components measured by Shannon’s index, two communities having identical indices can be differentiated by mapping richness and evenness coordinates on a scatter plot. In such graphs, confidence ellipses would allow testing significant differences between groups of samples. Multivariate statistical tests such as PERMANOVA can be performed on distance matrices calculated from richness and evenness coordinates and detect statistically significant differences that would have remained unforeseen otherwise.

## Introduction

Quantifying species diversity is a fundamental theme of ecology. Although there are several definitions of it (alpha, beta and gamma diversity), it is most often described in terms of alpha diversity, *i.e*. richness (the number of species) and evenness (a measure of how the species’ relative abundances tend to be uniformly distributed) within a community or habitat (Whittaker, 1960).

To summarize and compare the alpha diversity of two ecological communities, researchers frequently use scalar diversity indices. As they reduce the dimensionality of complex multivariate data into a scalar number, diversity indices can be compared using null hypothesis tests or confidence intervals (Hill, 1973). However, there is a myriad of those indices, each measuring different parameters, making the direct comparison of values from different indices difficult or even impossible. Some strictly measure species richness such as observed richness, Chao1 and ACE estimators (Willis, 2019) while others estimate alpha diversity as a phylogenetic metric (e.g. Faith’s PD index). Other metrics, such as Shannon’s index, englobe richness and evenness into a single metric. This index is unarguably one of the most popular metrics in community ecology, alongside Simpson’s diversity index, even though there is yet no clear guidelines on which diversity index should be used (Kim *et al*., 2017).

Shannon’s index (*H*) is defined as follows:

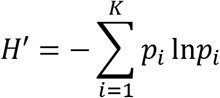

where *K* is the total amount of species in a biome and *p_i_* is the relative abundance (proportion) of species *i*. However, since it measures both richness and evenness in a single equation, there is theoretically an infinite number of richness / evenness value combinations translating into the same index score. Furthermore, richness and evenness may covariate positively, but also negatively. For example, in a 2020 study on microbiota dynamics in yellow perch (*Perca flavescens*) exposed to trace cadmium contamination, “decreasing richness and increasing evenness were observed” (Cheaib *et al*., 2020). There is no way of detecting whether evenness / richness covariance is either positive or negative by only using Shannon’s diversity index. To do so, Shannon’s index is usually compared alongside another index that measures either richness or evenness, e.g. Powell et al. (2015) or Sylvain *et al*. (2017).

Here is a contrived example with two simple mock communities to illustrate this issue (**Table 1**). Both have near-identical Shannon indices despite a very different community composition (1.609 and 1.608 respectively). This is because Shannon’s index binds richness 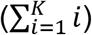 and evenness (−*p_i_*ln*p_i_*) together in a single equation. Decoupling its components would yield a more detailed overview of alpha diversity than what Shannon’s index would allow. This can be achieved by visualizing richness and evenness on 2D graphs, where each parameter would be assigned to an orthogonal axis.

**Table 1.** Relative species proportions in Communities 1 and 2.

| **Species** | **Proportion in**<br>**Community 1** | **Proportion in**<br>**Community 2** |
| --- | --- | --- |
| **Species 1** | 0.2 | 0.500 |
| **Species 2** | 0.2 | 0.100 |
| **Species 3** | 0.2 | 0.100 |
| **Species 4** | 0.2 | 0.100 |
| **Species 5** | 0.2 | 0.095 |
| **Species 6** | - | 0.050 |
| **Species 7** | - | 0.030 |
| **Species 8** | - | 0.025 |

## Methodology

### Deriving species richness

The simplest definition of species richness is the total amount of species found in a community. Richness can be derived from the summation operator in the Shannon index’s formula:

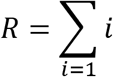

where *R* is species richness and *R* is the *i*-th species. Although several definitions of species richness have been formulated (e.g. Chao1, ACE, etc.), for the sake of simplicity, we will illustrate species richness herein by its simplest definition (i.e. the number of observed taxa without extrapolating rare taxa).

### Deriving evenness

Deriving evenness from Shannon’s index is not as obvious as deriving richness. The “evenness” component comes from the summation operand (−*p_i_*ln*p_i_*), where *p_i_* is the relative abundance of species *i*. The ln-transformation of *p_i_* in Shannon’s index’s formula narrows the range (and therefore the impact) of extreme values, and still weighs high relative abundances as “high”, and low abundances as “low”.

A way to estimate evenness would be to calculate the median of −*p_i_*ln*p_i_*. The median is an efficient trend indicator that is not affected by outlier values as the arithmetic average does (Welsh, 2004). Furthermore, a very uneven community would be expected to have a very low median −*p_i_*ln*p_i_*, whereas a perfectly even community would have a median −*p_i_*ln*p_i_* equal to any −*p_i_*ln*p_i_*.

As expected, evenness in Community 1 (0.321…) is higher than in Community 2 (0.227…). However, this formulation is not fully satisfying, as a perfectly even community (Community 1) should have an evenness index of 1, and a very unequal sample should have an evenness index nearing zero. In our example, indexes of even and uneven communities, although different, are very close. This issue could be solved by normalizing the median −*p_i_*ln*p_i_* by the highest −*p_i_*ln*p_i_* value. Normalized values of evenness become 1.000 for the perfectly even Community 1 and 0.655 for the uneven Community 2. To further validate this index, hereby called “normalized-median evenness”, let two additional communities be created, which are even more extreme than the first two, but each having three species in order to keep richness constant (**Table 2**). Normalized-median evenness clearly differentiates Communities 2 and 4, both of which are very different in terms of richness and evenness. Here is its definition:

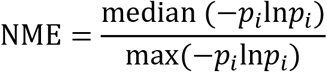

where NME is normalized-median evenness, *p_i_* is the relative abundance of species *i* and max is the maximum value of −*p_i_*ln*p_i_*.

**Table 2.**
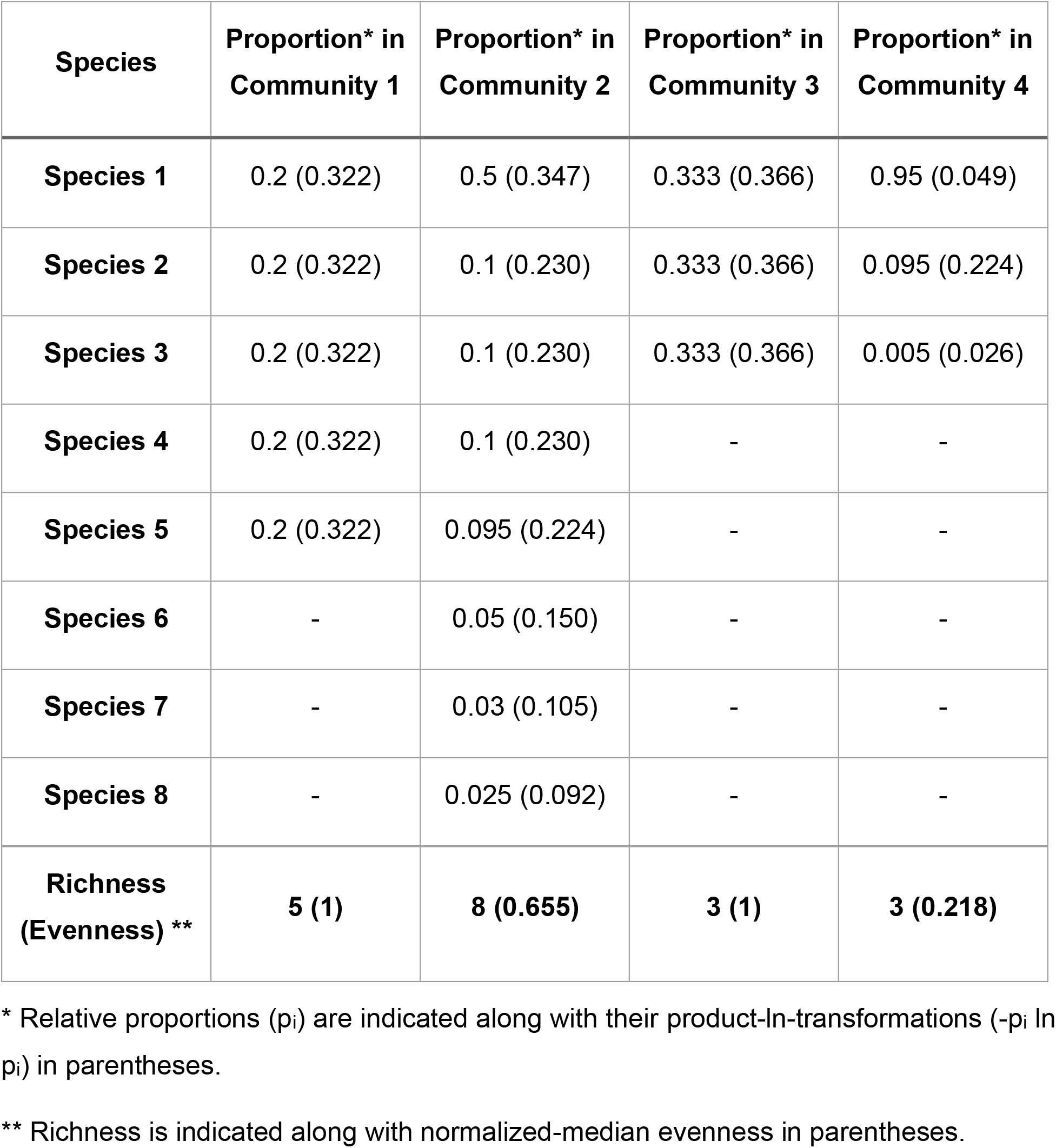
Relative species composition of mock communities 1 to 4 along with their product-ln-transformations.

### Graphical representation

By having both components of Shannon’s index deconvoluted, two samples can be compared simultaneously, even if they possess identical Shannon indices. This can be achieved by plotting richness and normalized-median evenness on a scatter plot, where each metric would correspond to a different axis (**Figure 1**). Communities 1 and 2 can be fully differentiated, even though they have the same Shannon index but different richness and evenness terms. Communities 3 (very even) and 4 (very uneven) are also fully differentiated from each other, even though their species richness is the same.

**Figure 1.**
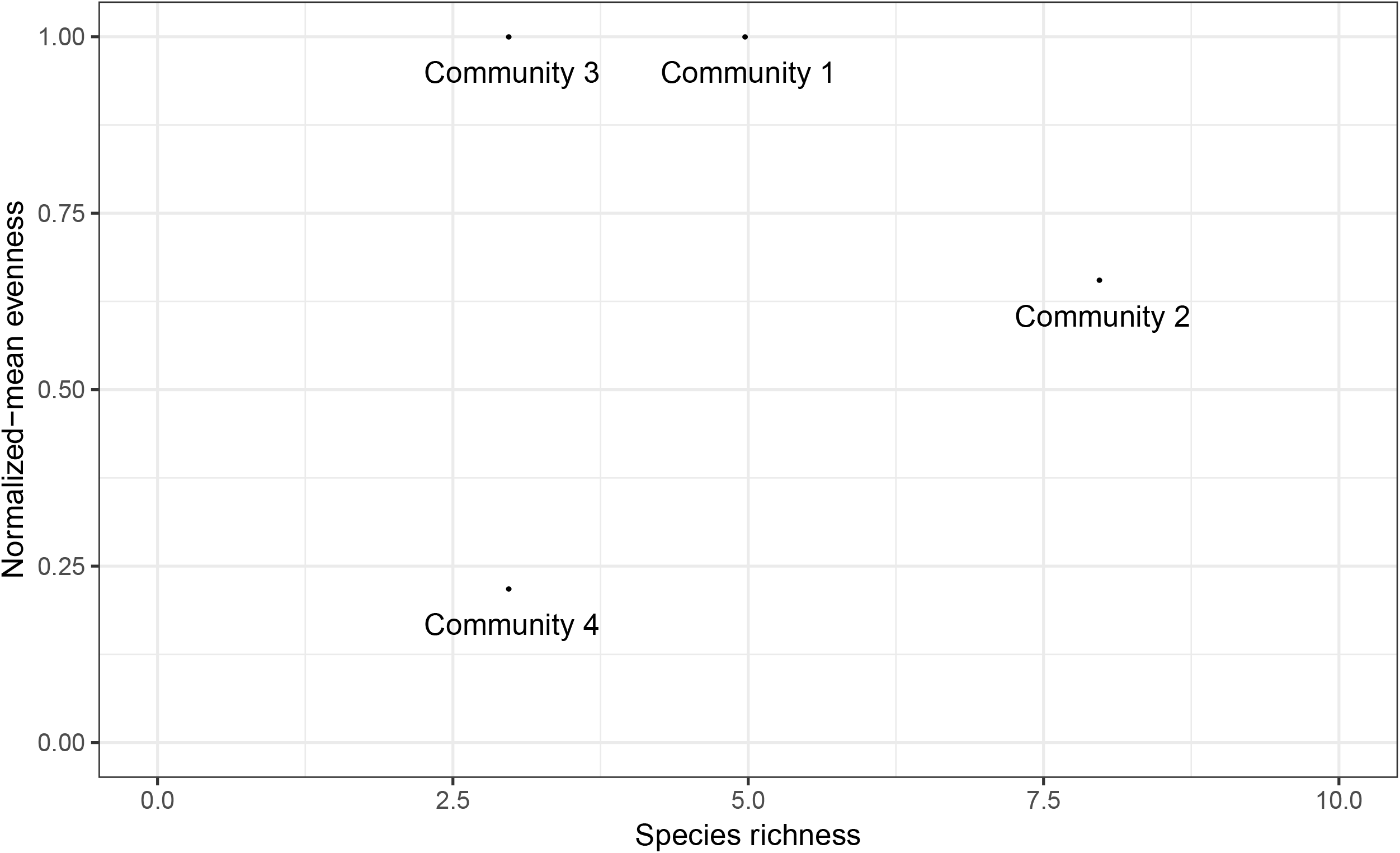
Normalized-Median Evenness vs Richness plot for mock communities 1 to 4.

### Example with a larger mock dataset

Let a mock dataset where the diversity of two groups (Alpha and Omega) of five samples (A to J) belong to two regions (Urban, Rural) is compared (**Table 3**). All previously discussed diversity indices, or their components, have been precomputed. Now let the data be plotted on a Richness vs Normalized-Median Evenness plot as previously described, with 95% confidence ellipses for each group (**Figure 2A**). A clear separation between samples from the Alpha and Omega groups can be made, which would not have been possible by comparing their Shannon indices alone (**Figure 2B**). Note how means and CIs overlap. Confidence ellipses may be used to detect statistically significant differences between groups, but they are not very useful for assessing the effect of several grouping variables at the same time. To illustrate this factor, let another grouping factor (Region) be included to fictional samples A to J mentioned above (**Table 3**).

**Table 3.**
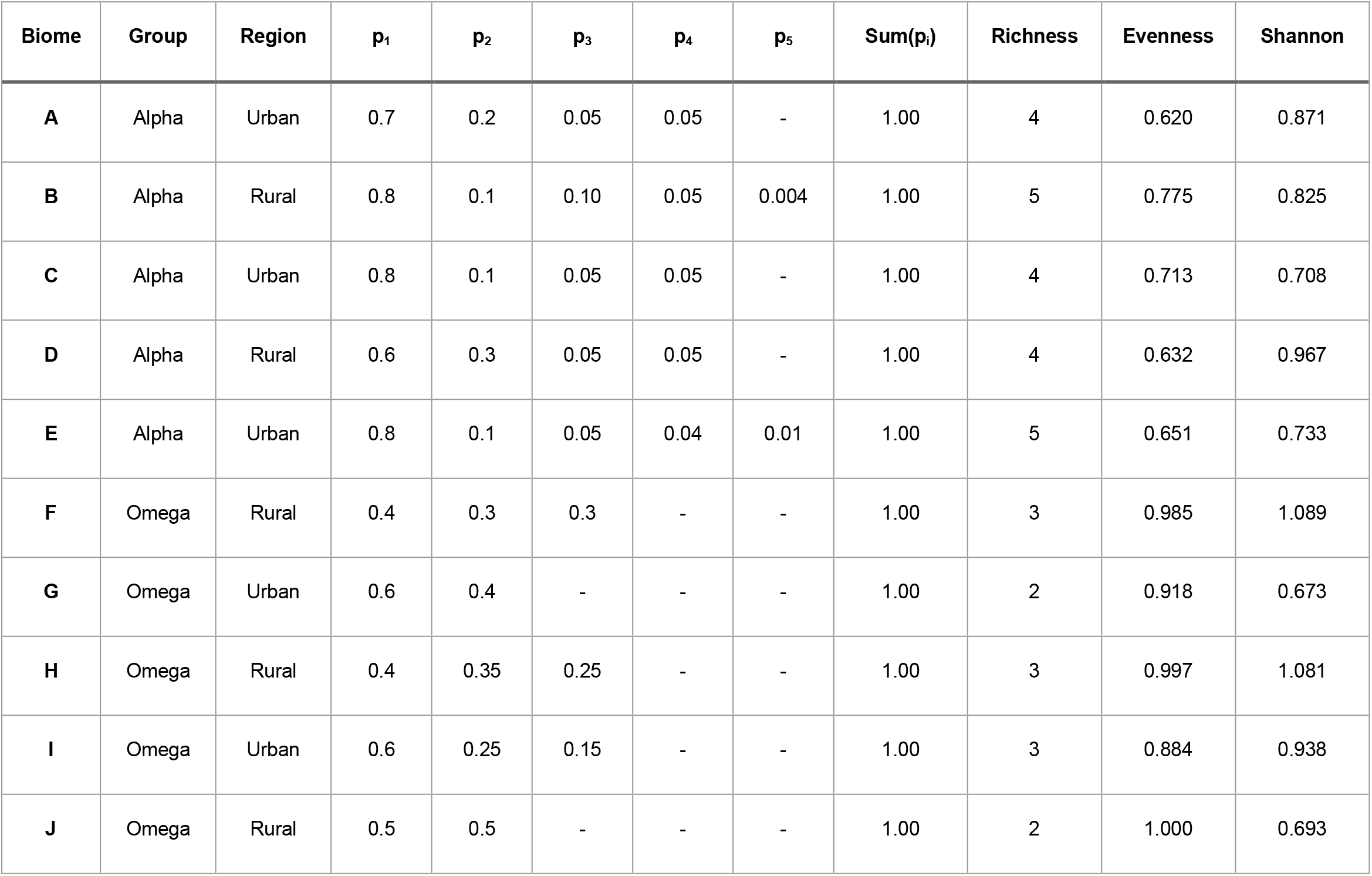
Species composition and various diversity metrics of a larger fictional dataset of 10 biomes belonging to two groups (Alpha and Omega) and two regions (Urban, Rural).

**Figure 2.**
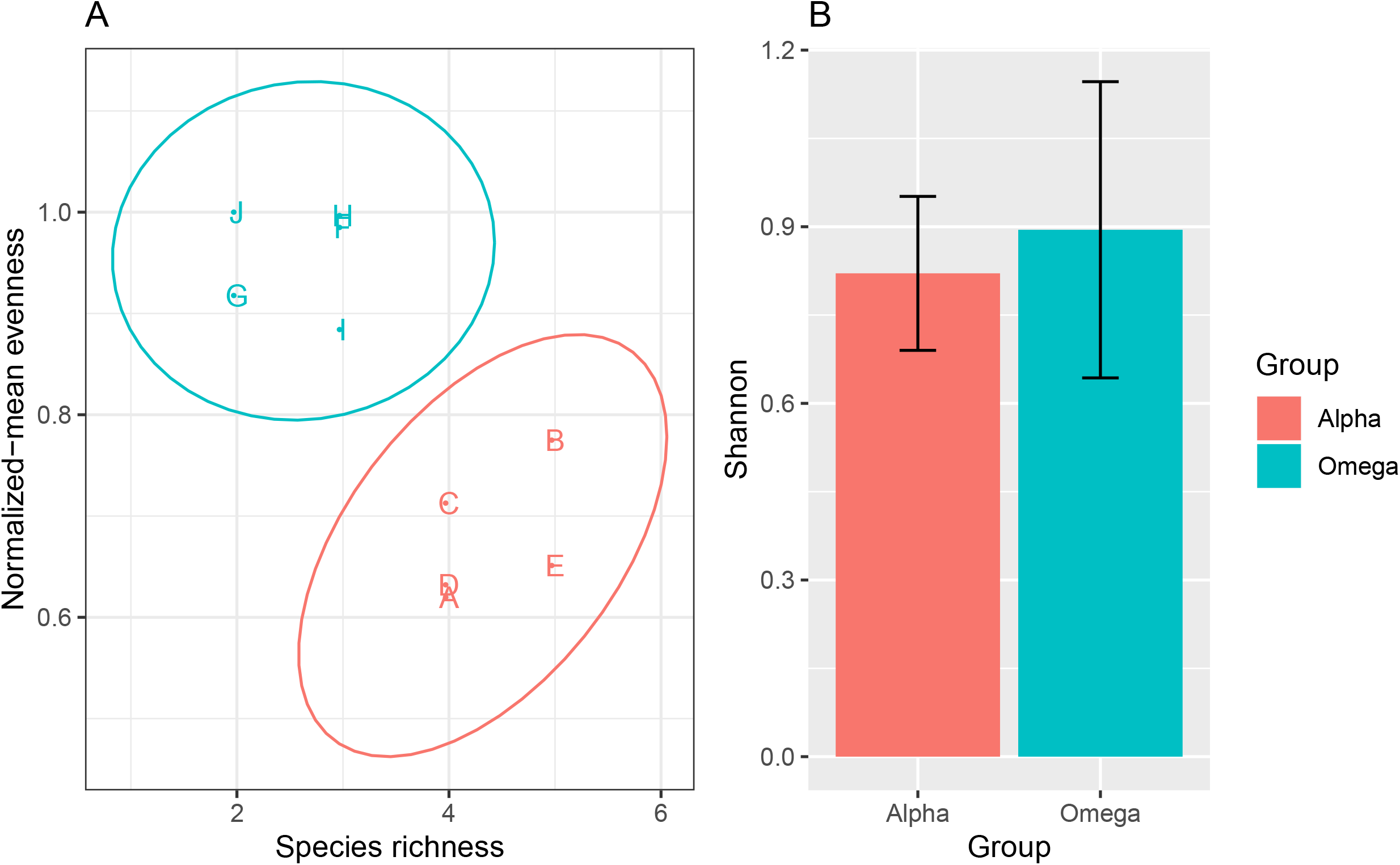
Alpha diversity for fictional communities A to J. (A) Normalized-Median Evenness vs Richness plot. Ellipses = 95% confidence intervals. (B) Mean Shannon index + 95% CI of Alpha vs Omega groups (n = 5 samples/group).

It is possible to use the evenness and richness values to compute an Euclidean distance matrix, which can be used as input for permutational multivariate analysis of variance (Anderson, 2001). A PERMANOVA was computed (99 permutations total) with the adonis() function from the vegan package (Dixon, 2003) using Richness and Evenness values as response variables and Group and Region as explanatory variables. It revealed a significant effect of Group on alpha-diversity (*F* = 14.3, *R*^2^ = 0.70, *P* = 0.02), but no significant Region effect (*F* = 0.32, *R*^2^ = 0.02, *P* = 0.59) and no significant interaction between Group and Region (*F* = 0.04, *R*^2^ = 0.0002, *P* = 0.88). If a conventional two-factor ANOVA had been performed using Shannon’s index as a response variable, no significant effect on alpha-diversity would have been detected, regardless of the grouping factor (0.2 < *P* < 0.92).

The next section details an example of real data analysis using Evenness-Richness scatter plots and related statistical analyses. Briefly, the use of those graphs allowed a more thorough view of alpha-diversity than using Shannon’s index. Moreover, we identified a clustering effect caused by the pooling of data obtained through multiple next-generation sequencing technologies. This effect was shown to be significant with PERMANOVA.

## Analysis of a real dataset: Enterotypes of the human gut microbiome (2011)

### Introduction to the dataset

Published in Nature in 2011, this work compared the fecal microbiota from 22 subjects using complete shotgun DNA sequencing (Arumugam et al., 2011). Authors further compared these microbial communities with the fecal communities of subjects from other studies. A total of 280 fecal samples / subjects are represented in this dataset, and 553 microbial genera were detected. The authors claim that the data naturally clumps into three community-level clusters, or “enterotypes”, that are not immediately explained by sequencing technology or demographic features of the subjects. This data is included into the R package phyloseq (McMurdie & Holmes, 2013) as an example dataset.

When studying the top ten most abundant genera across the enterotype dataset, we see that each enterotype is dominated by distinct subsets of genera (**Figure 3A**). Enterotype 1 is dominated by *Bacteroides* spp., whereas Enterotypes 2 and 3 are respectively dominated by *Prevotella* spp. and *Blautia* spp. Despite the very different top genera abundance profiles, the three enterotypes appear very similar in terms of alpha diversity when measured with Shannon’s index (**Figure 3B**).

**Figure 3.**
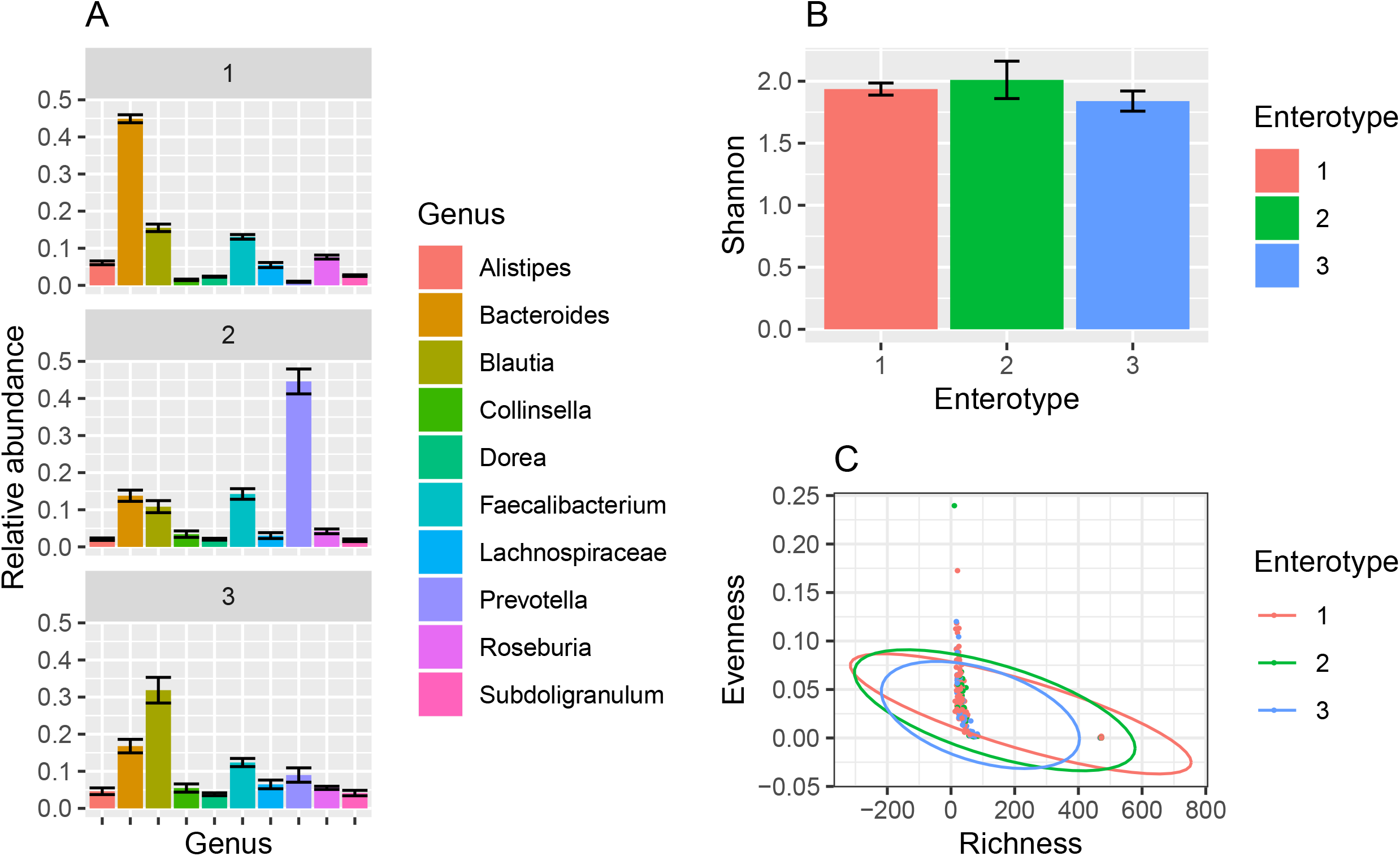
Impact of enterotype on alpha diversity within the human gut microbiota. (A) relative proportions of the top ten most abundant genera across the enterotype dataset. Error bars indicate 95% confidence intervals. (B) Mean Shannon index of enterotype samples grouped by enterotype. Error bars indicate 95% confidence intervals. (C) Evenness-Richness plot of enterotype samples grouped by enterotype. Ellipses = 95% confidence intervals.

### Evenness-Richness graph analysis

When alpha diversity across enterotypes is visualized with Evenness-Richness scatter plots instead, we see that the confidence ellipses of each enterotype group are entirely overlapping (**Figure 3C**). However, there are two completely distinct clusters visible on this figure, each being composed of samples from various enterotypes. In the right-side cluster (hereby named Cluster 2), there is a higher proportion of samples from Enterotype 1 (**Table 4**).

**Table 4.** Number of samples belonging to enterotypes 1, 2 and 3 in both clusters seen on the previous figure.

| Cluster | Samples from Enterotype 1 | Samples from Enterotype 2 | Samples from Enterotype 3 | Total |
| --- | --- | --- | --- | --- |
| Cluster 1 | 105 | 26 | 55 | 186 |
| Cluster 2 | 63 | 9 | 13 | 85 |
| Total | 168 | 35 | 68 | 271 |

There is a significant discrepancy on the relative proportion of enterotypes in both clusters (*χ*^2^ = 8.1945, *P* < 0.02). Given that those clusters are differentiated along the Richness axis, there appears to be a systematic bias on the assessment of richness between both clusters. Interestingly, the enterotype dataset includes data obtained through three different sequencing technologies (i.e. Sanger, 454 and Illumina). An Evenness vs Richness plot with samples labeled by sequencing technology revealed that Cluster 2 is made of all the Illumina samples of the dataset (**Figure 4A**).

**Figure 4.**
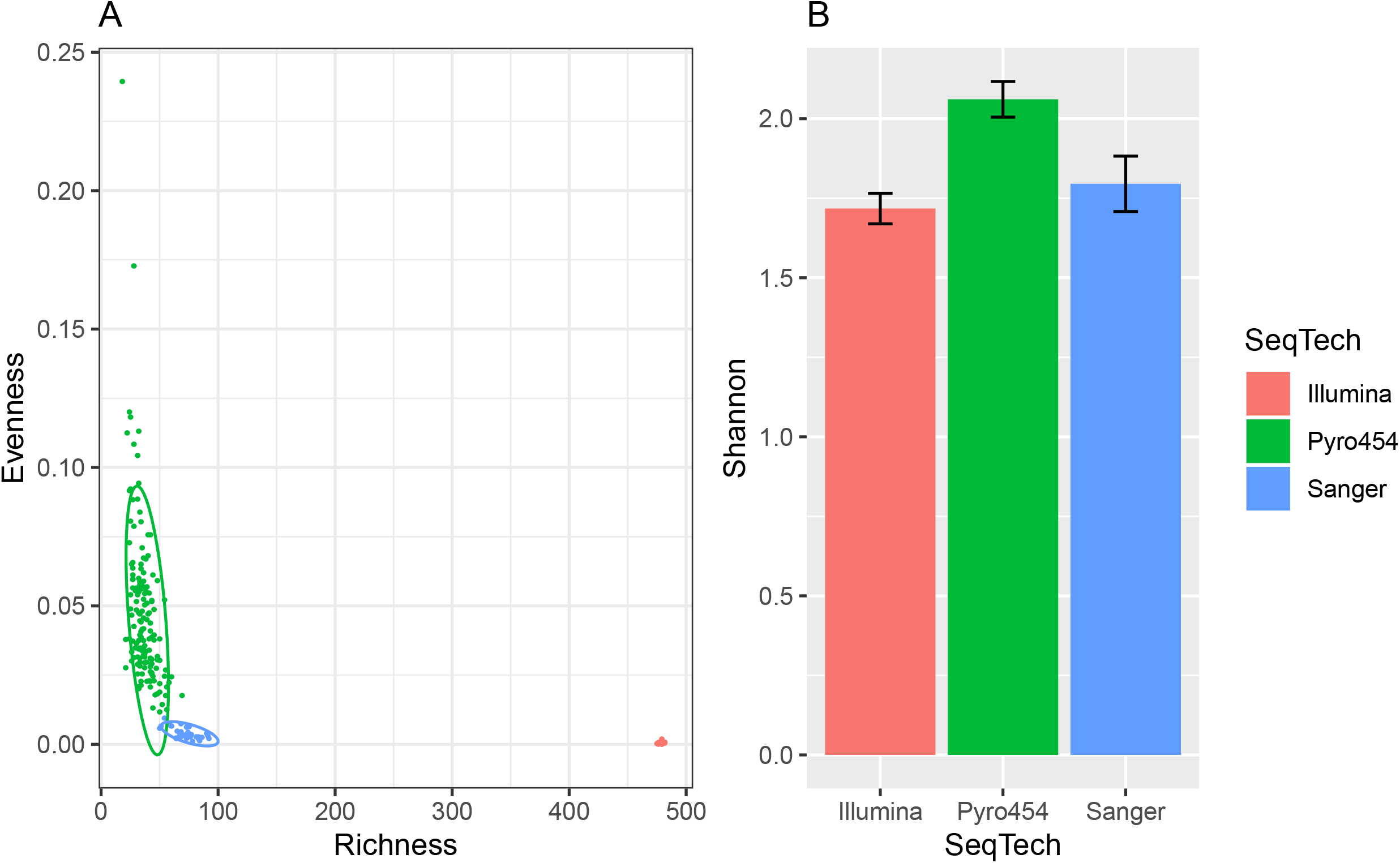
Alpha diversity of enterotype samples grouped by sequencing technology. (A) Evenness-Richness plot. Ellipses indicate a confidence level of 95%. (B) Mean Shannon index of enterotype samples grouped by sequencing technology. Error bars indicate 95% confidence intervals.

### Clustering effect caused by sequencing technology

Richness in Illumina samples is about one order of magnitude higher than in Sanger samples. This may be reflective of the high throughput of Illumina sequencing relatively to Sanger sequencing (Vincent et al., 2017). In contrast, evenness appears lower in Illumina samples than in Sanger and 454 samples. Those differences in either richness or evenness shed light on biases caused by sequencing technologies that could not have been detected while using Shannon’s index alone.

The average Shannon index of 454 samples is significantly different from the two other groups, whose means and CIs completely overlap (**Figure 4B**), despite that Illumina and Sanger samples differ in orders of magnitude in evenness. If the data from Cluster 1 (Sanger/454 samples) and Cluster 2 (Illumina samples) were analyzed separately, the conclusions made on the enterotypes’ alpha diversity would have been different (**Figure 5**).

**Figure 5.**
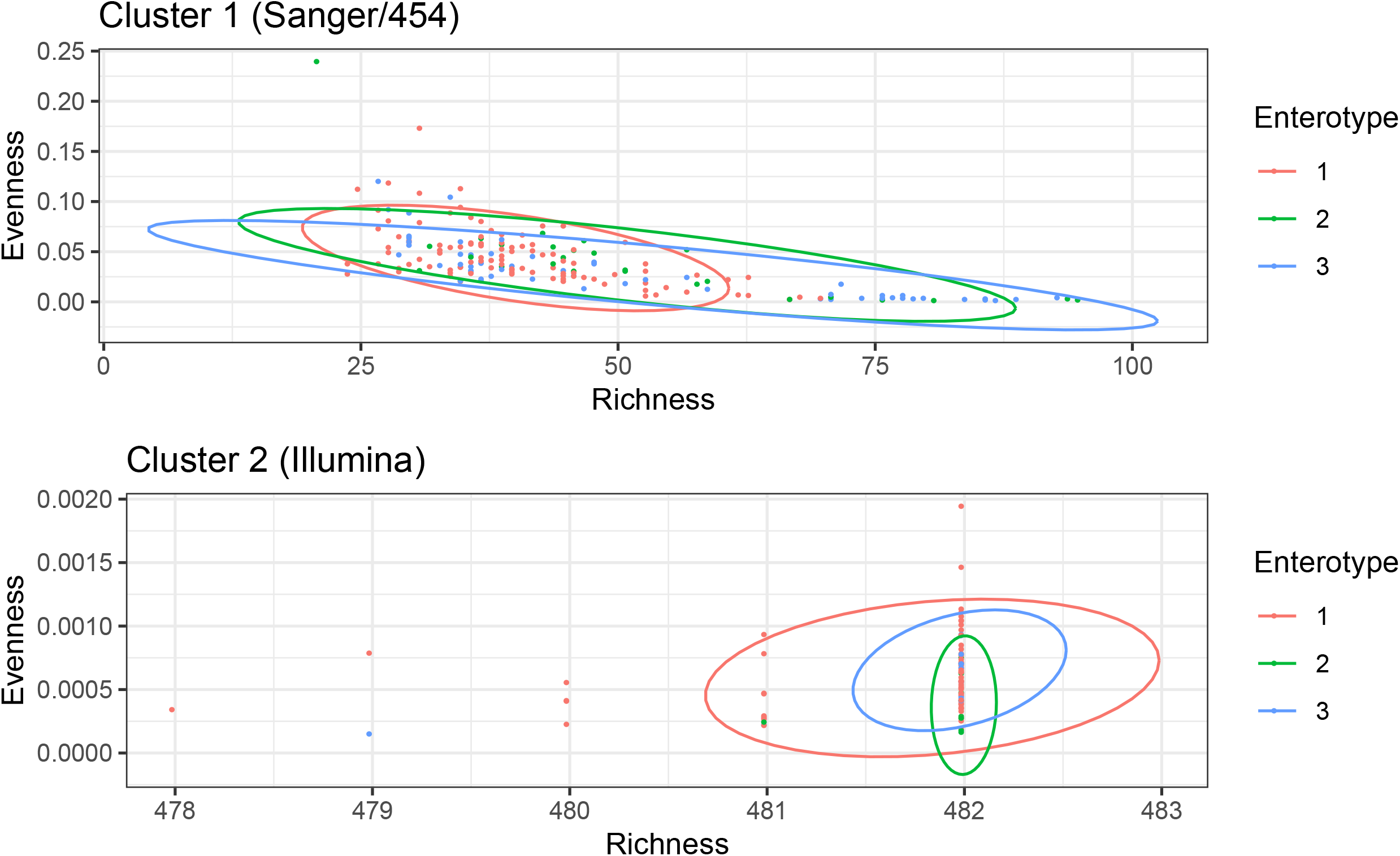
Evenness-Richness plot of samples in Clusters 1 (Sanger/454) and Cluster 2 (Illumina).

In Cluster 1 (Sanger/454), richness varies between ca. 25-100 genera and evenness between 0.01 and 0.25. Because the confidence ellipses of each enterotypes are not influenced by the outlying Illumina data, we can see that Enterotype 1 samples are more tightly clustered than enterotypes 2 and 3. This suggests a higher stability of alpha diversity with the gut microbiota of individuals with enterotypes 1, as opposed to individuals belonging to enterotypes 2 and 3.

In addition, richness in Illumina samples is 4-10 times superior to the richness of samples in Cluster 1 (Sanger/454). Knowing that Illumina sequencing technology has a maximum read throughput at least one order of magnitude above 454 pyrosequencing (Vincent et al., 2017), the differing richness values between samples of Clusters 1 (Sanger/454) and 2 (Illumina) may be attributed to the uneven sequencing depth between the three different technologies.

### Use of PERMANOVA to assess the clustering effect

The clustering effect caused by sequencing technology, and its influence on the distribution of enterotypes, were further assessed with a PERMANOVA. A total of 99 permutations were calculated, with the Evenness-Richness Euclidean distance matrix as a response object and Cluster and Enterotype as explanatory variables. There is a strong and significant effect of Cluster (*R*^2^ = 0.99, *P* < 0.01) and a small but significant effect of Enterotype (*R*^2^ = 0.00044, *P* < 0.01), on alpha-diversity, along with a small interaction effect between both factors (*R*^2^ = 0.0001, *P* < 0.03). Therefore, the probability that a sample belongs to a given enterotype is indeed mainly correlated with the sequencing technology used to generate the data.

### Influence of sequencing technology on top genera

Beyond alpha-diversity metrics, this sequencing technology bias also impacted the abundance of top genera in the dataset (**Figure 6**). Strong differences can be observed in the abundance of the top five genera. Most importantly, the top five genera differ between sequencing technologies. Only three genera out of five are consistently part of the top five: *Bacteroides, Faecalibacterium* and *Prevotella*. As for the other taxa, *Lachnospira* was completely undetected in samples sequenced by 454 technology. *Blautia* was only found in 454 samples. *Bifidobacterium* was detected only in Sanger samples. *Roseburia* was found only in 454 and Illumina samples.

**Figure 6.**
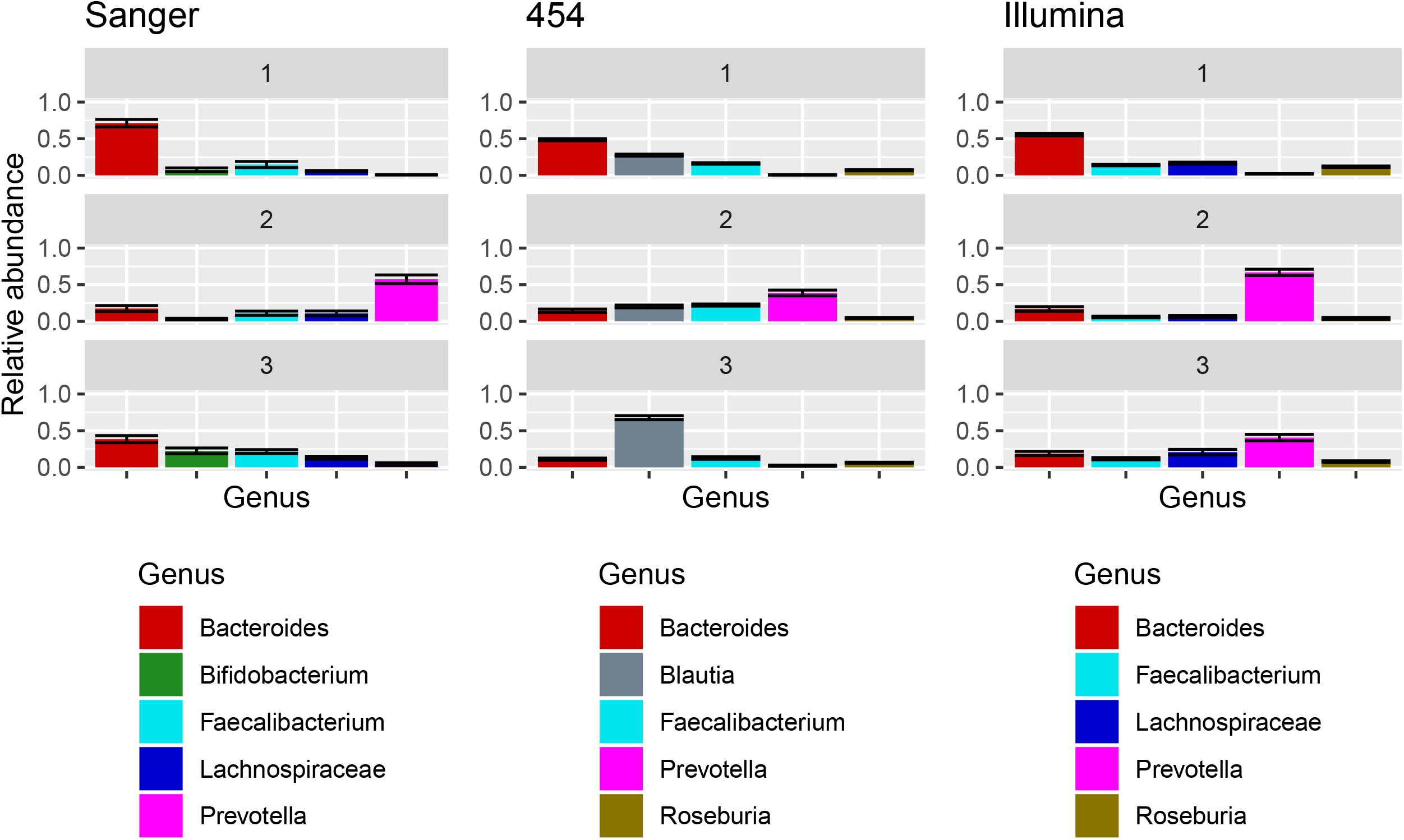
Impact of sequencing technology on the assessment of the top five most abundant genera across the enterotype dataset. Error bars indicate 95% confidence intervals.

We previously concluded that enterotypes 1 to 3 were dominated by *Bacteroides, Prevotella* and *Blautia* respectively (**Figure 6**). This observation holds true for enterotypes 1 and 2, but the dominance of *Blautia* in enterotype 3 is highly dependent on its high abundance in 454 samples (**Figure 6**). This highlights the challenges associated with merging microbiome datasets obtained through different sequencing methods in a single analysis. Notably, when technologies differ greatly in terms of sequencing throughput (e.g. Illumina vs. Sanger), richness estimates might be affected because the more a sequencing run produces reads, the more likely rare taxa are to be observed. Using an Evenness vs Richness plot to visualize alpha diversity allowed not only to detect but also to quantify this bias in the enterotype dataset.

## Conclusion

By comparing Shannon’s index alone, groups of samples may be entirely indistinguishable from one another. Moreover, one may overlook methodological biases that may affect the interpretation of alpha diversity analysis, e.g. combining datasets obtained though different sequencing technologies as seen in the enterotype dataset. Therefore, plotting the two components that it measures (richness and evenness) on 2D graphs gives a more thorough understanding of how alpha diversity differs between groups of samples. The data can be visualized in a 2D scatter plot where tight grouping indicates similarity between samples. Statistical methods, such as confidence ellipses or PERMANOVA, can be used to detect significant differences between groups, even if their Shannon index is the same.

## Materials and Methods

All statistical analyses were performed in RStudio using R v3.4.2. Briefly, mock datasets were prepared using pre-determined taxa abundance values in order to best illustrate cases where Shannon indices are identical between two communities despite different richness and evenness values. The enterotype dataset was imported from (and analyzed with) the phyloseq package suite for microbiome data analysis (McMurdie & Holmes, 2013). Top taxa abundance graphs were generated with a customized version of phyloseq’s plot_bar function which build mean taxa abundance plots with standard error of the mean (SEM) bars (see plot_bar_err.R in appendix). Shannon’s index and Evenness-Richness scatter plots were generated with package ggplot2 (Wickham, 2009) from summarized data remodeled from phyloseq-class objects with the summarySE function (see summarySE.R in appendix). PERMANOVAs were computed with the adonis() function from the R vegan package (Dixon, 2003).

## Data availability

A full R Markdown version of this manuscript (including code) is available on GitHub: http://www.github.com/jeffgauthier/alpha-diversity-graphs/.

## Acknowledgements

The authors wish to thank François-Étienne Sylvain and anonymous reviewers for critical insights and comments.

